# Comparative phenotypic, genomic and transcriptomic characterisation of two *Salmonella* Typhimurium strains for a first-in-human challenge model

**DOI:** 10.1101/2025.04.24.650425

**Authors:** Christopher Smith, Anan Bzami, Xiaojun Zhu, Jessica A. White, Nerie Roa, Akamol E. Suvarnapunya, Emma Smith, Anna Rydlova, Robert Varro, Zainab Kiliddar, Lijuan Luo, Blanca Perez Sepulveda, Melita A. Gordon, Graham S. Cooke, Robert K.M. Choy, Jay C.D. Hinton, Malick M. Gibani

**Affiliations:** Department of Infectious Disease, Imperial College London, London, UK; PATH, Seattle, WA, USA; Institute of Infection, Veterinary and Ecological Sciences, University of Liverpool, Liverpool, UK; Department of Diarrheal Disease Research, Walter Reed Army Institute of Research, Silver Spring, MD, USA; North-West London Pathology, Imperial College Healthcare NHS Trust, London, UK; Usher Institute, University of Edinburgh, Edinburgh, UK

**Keywords:** *Salmonella*, *Salmonella* Typhimurium, Non-typhoidal *Salmonella*, Human Challenge Model, Controlled Human Infection Model, Vaccines

## Abstract

**Background:** Invasive non-typhoidal *Salmonella* (iNTS) disease remains a major public health challenge in sub-Saharan Africa. *Salmonella* Typhimurium is responsible for the majority of cases, with specific lineages being associated with increased risk of bloodstream infection. We have recently developed a *Salmonella* Typhimurium controlled human infection model (CHIM) to better understand disease pathogenesis and to provide a platform to test candidate vaccines. Selecting appropriate challenge strainsis a central design consideration in developing challenge model protocols. We describe the rationale, manufacture, and detailed characterisation of the two *Salmonella* Typhimurium strains used in the first-in-human NTS CHIM.

**Methods:** Two Salmonella Typhimurium strains, 4/74 (ST19) and D23580 (ST313), were selected from the UK Health Security Agency National Collection of Type Cultures and manufactured under Good Manufacturing Practice (GMP) conditions. Challenge agent stocks underwent microbial limits testing, viability and stability assessments, and phenotypic characterisation including growth kinetics, motility, acid sensitivity, and antibiotic susceptibility. Whole-genome sequencing was performed to confirm genetic stability post-manufacture. The effect of pre-challenge handling conditions was assessed in saline and sodium bicarbonate buffers, and bacterial survival was evaluated under simulated gastric and intestinal conditions using a modified Rossett–Rice model. Transcriptomic profiling was undertaken to determine whether sodium bicarbonate exposure altered expression of key virulence genes.

**Results:** Both strains retained their expected phenotypic characteristics, including reduced motility and melibiose utilisation in D23580. GMP stocks remained pure, viable, and stable post-manufacture, with no unexpected genomic mutations detected. Both strains were susceptible to clinically relevant antibiotics used in the study. Survival was maintained in neutral and mildly acidic conditions, with significant reduction below pH 3.5. Stability was preserved for up to one hour in saline buffer and during simulated gastric transit in the Rossett-Rice model. Transcriptomic analysis showed no significant changes in *Salmonella* pathogenicity island-1 or 2, or flagellar gene expression following sodium bicarbonate exposure

**Conclusions:** These findings confirm that the *Salmonella* Typhimurium 4/74 and D23580 strains retained genomic and phenotypic integrity post-manufacture. The resulting challenge stocks provide the foundation for an ongoing NTS CHIM that aims to advance understanding of NTS and iNTS pathogenesis and support candidate vaccine testing.

## 1 Introduction

Invasive non-typhoidal *Salmonella* (iNTS) is a major cause of community-acquired bloodstream infection in sub-Saharan Africa (sSA) and is associated with significant morbidity and mortality [1]. iNTS disease was estimated to have caused 535,000 cases and 77,500 deaths in 2017 [2]. The greatest burden is observed in children under 5 years, with recognised risk factors including anaemia, malnutrition, and recent malaria infection [3]. Immunocompromised adults with HIV are also susceptible; however, incidence of HIV-associated infection has been declining with improved access to effective antiretroviral therapy [4].

*Salmonella* Typhimurium (STm) is the most commonly isolated serovar in iNTS bloodstream infections, followed by *Salmonella* Enteritidis(SEn)[5]. STm multi-locus sequence type (ST)313 has been identified as the dominant circulating pathovar in sSA and is associated with multi-drug resistance to amoxicillin, chloramphenicol and co-trimoxazole [5,6]. Additionally, the dominant circulating *S*. Typhimurium strains responsible for the majority of infections have been shown to have undergone evolutionary changes, characterised by genomic degradation and pseudogene formation, which are postulated to represent host adaptation towards an invasive phenotype [6]. The relative contributions of host susceptibility and host-adapted bacterial factors in iNTS are incompletely understood.

In addition to challenges posed by antimicrobial resistance (AMR), public health interventions to control iNTS disease burden are hampered by an incomplete understanding of transmission reservoirs and a lack of preventative vaccines [7]. Several promising vaccine candidates are in early-stage human development, including core and O-polysaccharide (COPS)/FliC conjugate vaccines and those utilising the Generalised Modules for Membrane Antigens (GMMA) platform [8–11]. The leading candidate vaccines are likely to progress as multivalent formulations targeting both STm and SEn, potentially alongside antigenic targets for typhoidal *Salmonella*.

Controlled human infection models (CHIMs) can provide unique insights into mechanisms of disease pathogenesis and elucidate immune correlates of protection [12]. In some circumstances, CHIMs can also provide clinically relevant platforms for candidate vaccine efficacy studies, supporting licensure decisions [13]. This has been particularly true for enteric pathogens, where CHIMs have played a central role in accelerating development of vaccines against *S.* Typhi and *V. cholerae* [14–16].

We have recently established a first-in-human NTS CHIM. The Challenge Non-typhoidal *Salmonella* (CHANTS) model aims to characterise the clinical and microbiological responses of healthy volunteers to oral challenge with *S.* Typhimurium, describe putative immune correlates of protection, and provide a platform for future NTS vaccine efficacy assessments. A consortium of experts in the fields of *Salmonella* biology, epidemiology, vaccine development and human challenge models have contributed to the study design [17].

Several considerations relating to bacterial challenge strain selection, manufacture and characterisation are central to the safe and ethical conduct of controlled human infection models. The approach to strain selection has been described in detail for other bacterial human challenge models [18,19]. Whilst some pathogen-specific considerations may apply, some general principles are outlined in **Table 1**. To maximise the utility of a disease model, challenge strains should aim to be as a minimum: i) phylogenetically representative of contemporary disease-causing isolates; ii) susceptible to commonly used antibiotics; iii) have a traceable history from isolation; iv) possess a representative repertoire of virulence factors and v) be associated with a relevant disease phenotype for vaccine efficacy assessment. Here we describe the strategy used to select the most relevant strains for use in a novel NTS CHIM. We provide comprehensive phenotypic and genomic characterisation of selected strains and detail the testing protocols associated with manufacture to GMP standards. We go on to explore the effect of sodium bicarbonate exposure on *S.* Typhimurium virulence gene expression, in an attempt to model the bacterial transcriptional profile at the point of oral administration.

**Table 1:** Key considerations for challenge agent selection, development and production for controlled human infection models. CA: challenge agent, CHIM: controlled human infection model, QC: quality control. Adapted from [20].

| Consideration | Rationale |
| --- | --- |
| Challenge Agent Strain Selection | • Representative of current strains causing natural infection or disease by the pathogen under investigation<br>• Selected to produce measurable clinical symptoms or signs which mimic natural infection with the pathogen under investigation (established by pre-determined clinical endpoints), without causing severe disease or complications.<br>• Wild type, attenuated, or genetically modified strains may be utilised, determined by the primary objective of the CHIM and its intended application e.g. vaccine efficacy assessment.<br>• Full provenance of the challenge agent must be known and should have a traceable history from isolation |
| Challenge Agent Strain Characteristics | • Reliable susceptibility to antibiotics intended for use within the context of a CHIM<br>• Demonstrated genetic stability throughout the study period to avoid unexpected changes in virulence or resistance |
| Challenge Agent Strain Manufacture, QC and Testing | • Equipment and reagents must be available, serviced, calibrated, and operated by trained personnel<br>• Production should be within laboratory facilities with appropriate Biosafety containment level suitable for the specific CA<br>• Key physical and chemical properties of the CA must be demonstrated and recorded: identification via standard microbiological diagnostics, absence of impurities or toxins, stability and sensitivity to pH, temperature chemicals<br>• Purity: CA should be free from other agents of human infection (viruses, bacteria, fungi), determined by microbial limits testing<br>• Stability: CA should maintain stability following manufacture, transport and storage, determined by repeat testing and appropriate intervals |
| Regulatory Compliance | • CA dossier must be produced and available following release from manufacture, detailing testing results from the above listed requirements<br>• Engagement with regulatory bodies overseeing manufacture and production is essential and comply with local/national regulations e.g. international variation in classification of CAs as investigational medical products<br>• Full consideration must be given to public health and infection control aspects of CA use e.g. informing local health protection teams and developing risk assessments to manage unintentional release or transmission of CA<br>• Risk assessments must be in place for the import, transport and storage of CA within appropriate biosafety facilities |

## 2 Materials and Methods

### 2.1 CHANTS Study Design

The CHANTS study detailed protocol has previously been described [21]. In brief, healthy UK participants aged 18-50 are randomised to receive oral challenge with one of two strains of *S.* Typhimurium - D23580 (ST313) or 4/74 (ST19) – in a double-blind, 1:1 randomisation protocol. A dose-escalation protocol is applied using the continual-reassessment method [22]. The primary objective is to determine the minimum infectious dose in colony forming units (CFU) for 60-75% of participants to develop systemic Salmonellosis, defined as bacteraemia and/or sustained fever (oral temperature ≥38°C for ≥12 hours). Secondary objectives are to describe and compare the clinical, microbiological and immunological responses following challenge. Participants are monitored in a hospital environment for 7 days following challenge and are followed for 1 year to assess long-term safety.

### 2.2 Challenge Agent Strain Selection and Manufacture

STm D23580 and 4/74 stocks were sourced from the UK Health Security Agency National Collection of Type Cultures repository. Stocks were transferred to the Pilot Bioproduction Facility at Walter Reed Army Institute of Research, MD, USA for manufacture under Good Manufacturing Practice (GMP) conditions. Each challenge agent was prepared by inoculating a single colony of *Salmonella* Typhimurium from a research cell bank ampoule onto Tryptic Soy Agar (TSA) plates. A single selected colony was used to inoculate APS LB media in a baffled shake flask, which was incubated at 37 ± 1°C with shaking (200 ± 10 RPM) until the culture reached an OD600 of 2.0 ± 0.2. The culture was then mixed with 16% synthetic glycerol as a cryoprotectant, aliquoted into sterile cryovials, and flash-frozen for storage at ≤-70°C in ultra-low temperature freezers. Purity was ensured by streak plating, Gram staining, and dynamic environmental monitoring compliant with GMP standards.

Colony counting was performed to assess the pre- and post-freeze viability of the challenge strains. Two vials of each strain were thawed, and serial dilutions (10^-1^ to 10^-7^) were prepared in 0.9% saline. Triplicate TSA plates were inoculated with 100µL of the 10^-5^ to 10^-7^ dilutions and incubated at 37±1°C for 18-20 hours. Colony counts were conducted on plates containing 30– 300 colonies. Agglutination testing was conducted using the Sero-Quick Group kit to confirm serogroup B antigen expression. Colonies were transferred to drops of Group B antiserum or 0.9% saline on a glass slide and observed for visible agglutination within 10 seconds.

### 2.3 Microbial Limits Testing

Manufactured challenge agent stocks underwent microbial limits testing for the presence of non-sterile products, including specified organisms: Bile Tolerant Gram-Negative bacteria, *Escherichia coli, Pseudomonas aeruginosa, Staphylococcus aureus, Bacillus spp., Clostridia spp., Aspergillus spp.,* and *Candida spp*. Stocks were diluted in TSB diluent 10^-3^ to 10^-8^ and plated on a range of organism specific selective media for microbial enumeration.

### 2.4 Acid Sensitivity of *S.* Typhimurium in low pH buffers

A pH range from pH 5 to pH 3.5 was used to evaluate acid sensitivity of each challenge strain. Test bufferswere prepared in 50ml conical tubes with a mixture of simulated gastric fluid (SGF; 7108-32, Lot 2301H18), sodium bicarbonate (Fisher S233-500, Lot 220180), and 0.9% sterile saline (KD Medical RGF-3290, Lot 102021-0) to achieve the desired test pH. Cells were added to each of the pH buffers at a concentration of 5.15x10^5^ CFU/ml and held in a 37°C water bath for pH adjusted samples or on ice for control samples for 15 minutes. After incubation the buffers were neutralized with sodium bicarbonate and diluted with 0.9% saline to a final volume of 32.1ml. Each test sample wasplated in triplicate on LB(Teknova, L7590) media (0.1ml per plate) and incubated overnight at 37°C prior to measuring mean CFU counts (n=3).

### 2.5 *S.* Typhimurium Stability in 0.9% Saline Compared with Sodium Bicarbonate Buffer

Stability of holding the challenge agents in bicarbonate buffer (2.6g/120ml) or 0.9% saline solution was evaluated first with D23580. The challenge agent was diluted in saline or bicarbonate buffer to 5.15x10^5^ CFU/ml and held on ice for up to 3 hours. 0.1 mL of solution was sampled at hourly intervals and plated on LB prior to overnight incubation at 37°C. Mean CFU counts were determined from three replicate LB plates per time point, with the experiment being performed in duplicate. Upon identification of optimal pre-challenge holding conditions for D23580 (0.9% saline; 1-hour hold), the relevant conditions were tested and confirmed with 4/74 by diluting in 0.9% saline and holding for 1 hour on ice prior to plating and counting as described above.

### 2.6 Simulated Gastrointestinal pH Dynamics in a Modified Rossett-Rice Model

Stability of the challenge strains was evaluated using a modified Rossett-Rice model (RR) at a 10-fold reduced scale [23]. A sodium bicarbonate dose of 0.26g (representing a 2.6g human dose) was evaluated and represents the formulation used in previous *Salmonella* Typhi and Paratyphi challenge models [24].

To first determine the pH range expected in the RR model, a peristaltic pump (set at a rate of 0.4 ml per minute) was used to add additional SGF throughout the experiment to mimic conditions in the stomach. Additionally, a 1000ml beaker water bath was prepared and set to 37°C with a 200ml beaker placed inside. The stomach contents comprised 0.26g sodium bicarbonate in 12ml sterile water for injection (WFI; KD Medical RGF-3410, Lot 081720-04), 3ml of sterile 0.9% saline, 0.1ml of 0.9% saline (substituted for cells), and 5ml of simulated gastric fluid (SGF; 7108-32, Lot 2301H18). This resulted in a final buffer dose of 0.26g. The contents of the 50ml conical tube were poured into the 200ml beaker and temperature and pH were monitored every minute for a total of 70 minutes after an initial 5 minutes for acclimation.

### 2.7 *S.* Typhimurium Stability in a Modified Rossett-Rice Model

To evaluate challenge strain stability in the RR model, the RR was repeated with 0.1 ml of cells diluted in 0.9% saline or 0.26g bicarbonate buffer to a working concentration of 5.15x10^5^ CFU/ml. Cells were held in 50ml conical tubes on ice for zero, one, and two hours prior to testing in the Rossett-Rice model. The ability to recover bacteria was determined by plating at 0, 30, 45, and 60 minutes. Time points were plated on four replicate LB plates (0.1ml per plate) and incubated overnight at 37°C prior to CFU counting.

To evaluate the challenge strain stability after transit into the small intestine, the RR model was conducted as described above for 70 minutes, before samples were removed and added to simulated intestinal fluid (Ricca 7109-32, Lot # 1412H21). 3.88ml samples of the RR (“gastric fluid”) were taken at 0, 30, 70min and mixed with 26.1ml of simulated intestinal fluid for a gastric fluid to intestinal fluid ratio of 31cm^3^/209cm^3^ [25]. Simulated intestinal fluid was filtered prior to use and heated to 37°C. After addition of the gastric fluid to the simulated intestinal fluid the solution was held at 37°C with stirring at 65 RPM. Samples of 0.1ml were plated and counted as outlined above following overnight incubation.

### 2.8 Phenotypic Characterisation

#### 2.8.1 Growth Characteristics

Growth curves for each strain were determined by culture in the Enzyscreen Growth Profiler in a range of nutrient conditions. LB overnight cultures were diluted to optical density OD_600_ = 0.01 for inoculation into 96-well plates. Plates were incubated at 37°C for 24 hours with shaking at 220rpm, and measurements obtained every 30 minutes. The Growth Profiler converts green pixel values of each well to optical density based on established calibration curves for relevant media. Each growth curve was generated by the mean values of six replicates. Growth curves for D23580 and 4/74 were assessed in LB media, M9 minimal media supplemented with 5% glucose, and M9 minimal media supplemented with melibiose.

#### 2.8.2 Swimming Motility

Differential swimming motility between strains was assessed by spotting 3µl LB overnight culture of D23580 and 4/74 onto 0.3% LB agar. Plates were left at room temperature for 30 minutes prior to incubation at 37°C. The diameter of migration halos was assessed at 4 hours to determine the relative bacterial swimming motility between strains.

#### 2.8.3 Red, Dry and Rough (RDAR) Morphotype

Red, dry and rough (RDAR) morphotype was assessed in NaCl-deficient LB plates supplemented with 40µl/ml Congo red. 2µl of LB overnight culture was spotted onto plates and incubated at room temperature for 72 hours prior to imaging colonial morphology using the ImageQuant 4000 biomolecular imager.

#### 2.8.4 Antibiotic Susceptibility

Antibiotic susceptibility testing was undertaken by disc diffusion and broth microdilution in accordance with EUCAST guidelines [26]. Disc diffusion was performed on Mueller-Hinton (MH) agar with antibiotic discs applied and cultured overnight at 37°C. Interpretation as sensitive (S) or resistant (R) was determined from zone diameter breakpoints produced by EUCAST. Antibiotics tested include ampicillin (10µg), pefloxacin (5 µg), chloramphenicol (30 µg), trimethoprim-sulphamethoxazole (25 µg), ciprofloxacin (5 µg), and ceftriaxone (30 µg).

Broth microdilution was undertaken using the automated BD Phoenix system. Minimum inhibitory concentrations were measured in accordance with EUCAST guidelines to determine susceptibility [27]. All antibiotic susceptibility testing was performed in accredited diagnostic facilities at North-West London Pathology, Imperial College Healthcare NHS Trust.

### 2.9 Whole Genome Sequencing

Genomic DNA was extracted using the ZYMO Research Quick-DNA Miniprep Plus Kit according to the manufacturer’s instructions. In summary, 1ml overnight LB culture was lysed then incubated in a heat block at 55°C for 10 minutes. 420µl genomic binding buffer was added prior to centrifugation and repeated wash with DNA buffer. 30µl DEPC water was added prior to final centrifugation with resulting genomic DNA stored at -20°C. Illumina sequencing of extracted genomic DNA was undertaken (SeqCenter, Pittsburgh, PA, USA), and sequence data was analysed for mutations using SNIPPY software v4.6.0.

### 2.10 Investigation of Bicarbonate Exposure on *S.* Typhimurium Virulence Factor Gene Expression

The impact of sodium bicarbonate exposure on virulence gene expression by D23580 and 4/74 was evaluated by transcriptomic profiling. RNA was extracted from D23580 and 4/74 after incubation in either 0-minutes or 30-minutes incubation in 0.53g/30ml sodium bicarbonate solution. RNA was extracted from STm strains using a modified TRIzol-based protocol. Overnight cultures were grown in LB(10 g/L tryptone, 5 g/L yeast extract, 5 g/L NaCl) at 37°C with shaking (220 rpm). Cultures were diluted 1:1000 into fresh LB, grown to the desired OD_600_ = 2.0. Cells were centrifuged then mixed with stop solution (5% phenol [pH 4.3], 95% ethanol) to stabilize RNA either immediately (0-minutes) or after 30-minutes incubation in sodium bicarbonate solution. Samples were centrifuged at 3,220g for 10 minutes at 4°C, and bacterial pellets were resuspended and transferred to microfuge tubes. The pellets were lysed in 1ml TRIzol reagent on ice, and RNA was extracted using phase lock gel tubes. Following the addition of 400µl chloroform, samples were mixed, incubated, and centrifuged to separate phases. The aqueous phase was recovered, and RNA was precipitated with isopropanol. The resulting RNA pellets were washed with 70% ethanol, air-dried, and dissolved in 20µL DEPC-treated, RNase-free water. RNA quality and concentration were assessed using a Qubit fluorometer and Bioanalyzer (Agilent RNA 6000 Nano kit). RNA samples were treated with DNase using the Invitrogen RNase-Free DNase Kit to remove residual DNA. Libraries were prepared using Illumina’s Stranded Total RNA Prep Ligation with Ribo-Zero Plus kit, incorporating 10bp unique dual indices (UDI). Sequencing was performed on a NovaSeq X Plus platform, generating paired end 150bp reads. Demultiplexing, quality control, and adapter trimming were conducted using *bcl-convert* (v4.2.4). Sequence data was analysed in transcripts per million (TPM) to assess relative gene expression.

## 3 Results

### 3.1 Challenge Strain Selection

Strain selection for use in the CHANTS study has been conducted in accordance with the principles and considerations outlined in **Table 1**. The CHANTS study usestwodistinct strains of *S*. Typhimurium which are considered representative of both invasive (iNTS) and diarrhoeal (dNTS) disease-causing isolates. D23580 belongs to the dominant ST313 sequence type causing invasive disease in sSA, whilst 4/74 (ST19) is considered an archetypal strain representative of globally distributed infections causing enterocolitis [5]. In addition to their epidemiological relevance, D23580 and 4/74 are model organisms which have a traceable history from isolation, and have undergone extensive prior *in vitro* phenotypic, genomic, and transcriptomic characterisation, which facilitates a deeper understanding of biological properties [28,29]. Both strains have been deposited in the UK Health Security Agency National Collection of Type Cultures and have been sourced from this repository for the purposes of this study.

The final selection of these specific strains was informed by expert consultation and ultimately based on their status as well-characterised, archetypal laboratory strains representing the two major lineages of *Salmonella* Typhimurium [17]. These choices enabled us to leverage extensive microbiological, genomic and pathogenesis data, allowing presentation of strain characteristics – including virulence determinants, antimicrobial susceptibility, and epidemiological relevance – to regulatory trial sponsors and research ethics committees. Both strains have been extensively studied *in vitro* and *in vivo* and have served as references in numerous studies of *Salmonella* pathogenesis and host interactions, providing a strong evidence base to support their safety and suitability for use in a controlled human infection model (CHIM).

### 3.2 GMP manufacture, quantification, and purity of challenge stocks

Identification of manufactured challenge agent stocks was initially confirmed by positive agglutination of both strains with *Salmonella* group B antiserum. Pre- and post-freeze viability was confirmed by colony counting following serial dilutions. The pre-freeze viability of 4/74 was 1.24 x 10^9^ CFU/ml, and the post-freeze viability was 5.2 x 10^8^ CFU/ml. The pre-freeze viability of D23580 was 9.95 x 10^8^ CFU/ml, and the post-freeze viability was 4.9 x 10^8^ CFU/ml.

Microbial limits testing confirmed manufactured challenge agent stocks were free from non-sterile products, including specified organisms: Bile Tolerant Gram-Negative bacteria, *Escherichia coli, Pseudomonas aeruginosa, Staphylococcus aureus, Bacillus spp., Clostridia spp., Aspergillus spp.*, and *Candida spp*. Stocks diluted in TSB diluent 10^-3^ to 10^-8^ and plated on a range of organism specific selective media confirmed absence of objectionable organisms.

### 3.3 *S.* Typhimurium viability is reduced in acidic pH <3.5

We next evaluated the optimal conditions for challenge agent viability and delivery, focusing on the role of gastric acid pH as a first line of defence against enteric pathogens. Previous *S.* Typhi challenge studies demonstrated that neutralising gastric acid prior to administration significantly lowers the infectious ID_50_ dose [24]. Pre-treatment with sodium bicarbonate – routinely used in human challenge models for *S.* Typhi [24], *Vibrio cholerae*[30], and *Shigella* spp. [31] is also thought to reduce batch-to-batch variability and enhance reproducibility in enteric challenge studies.

We began by assessing the pH sensitivity of D23580 and 4/74, with the aim of defining a threshold below which viability of the challenge strains significantly decreases. 5.15x10^5^ CFU/ml of each strain were incubated for 15 minutes in several pH test conditions prior to plating. Both strains demonstrated similar stability in 0.9% saline, sodium bicarbonate solution (2.6g/120ml), and acidic buffer pH 5. Survival declined following incubation in pH ≤ 3.5, with both 4/74 and D23580 demonstrating a trend towards lower survival compared to other conditions **(Figure 1 – Panels A-B).** These findings are consistent with known sensitivity of *Salmonella*to acidic conditionsand supports the commonly utilised approach of neutralising gastric acid with bicarbonate pre-treatment in enteric challenge models [32,33].

**Figure 1.**
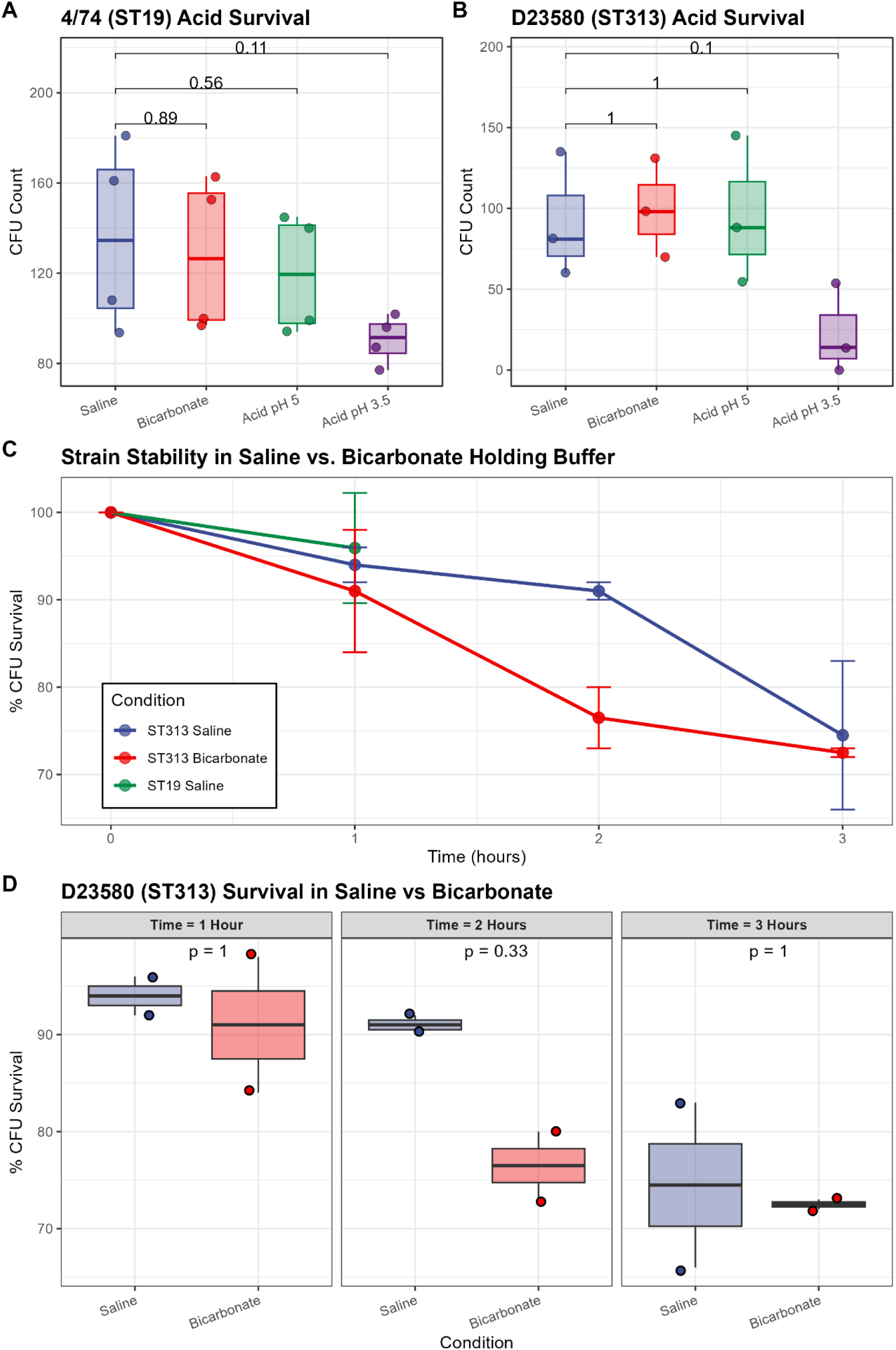
Acid pH sensitivity and stability of 4/74 and D23580 in saline vs. sodium bicarbonate holding buffers. (A) *S.* Typhimurium 4/74 and (B) D23580 survival following incubation for 15 minutes in 0.9% saline, 2.6g/120ml sodium bicarbonate, and acidic buffer pH 5 and pH 3.5. 0.9% saline served as the control buffer for pairwise comparisons. Boxplots represent median and interquartile range. (C) D23580 and 4/74 survival in 0.9% saline compared with 2.6g/120ml sodium bicarbonate buffer over time. Percentage CFU survival compared as a proportion with T=0hrs. D23580 was tested across all time points and buffer conditions. 4/74 was tested only in saline at T=0 and T=1hr following selection of the optimal conditions. (D) D23580 survival contrasting 0.9% saline with bicarbonate across timepoints. Boxplots represent median and interquartile range. P = two-sided Mann-Whitney U test.

### 3.4 *S.* Typhimurium viability remains stable in 0.9% saline and bicarbonate buffer to 1 hour

We sought to determine the stability of challenge strains under varying holding buffer conditions. We began by comparing bacterial survival following incubation in 0.9% saline and sodium bicarbonate (2.6g/120ml) buffer, in addition to testing different holding times on ice (T = 0, 1, 2, 3 hours) to determine the optimal solution and maximum hold duration to ensure viability at the time of oral administration. Testing was initially performed with D23580, and the relevant optimal holding conditions (0.9% saline; 1-hour hold) were repeated and confirmed for 4/74. Survival was maintained at >90% in 0.9% saline solution to T=2hrs, dropping to <80% after T=3hrs, compared with T=0hr baseline. There was a trend towards reduced survival in sodium bicarbonate compared with 0.9% saline, with a marked drop in survival to <80% at T=2hrs **(Figure 1 – Panels C-D)**. Similar survival >90% at T=1hr was demonstrated for strain 4/74. As a result of these findings, we amended the challenge agent preparation SOP to use 0.9% saline solution instead of sodium bicarbonate as the holding buffer with a maximum one-hour hold interval between formulation and administration.

### 3.5 Sodium bicarbonate pre-treatment maintains simulated gastric fluid pH above critical thresholds for STm survival

In the CHANTS study, participants are administered a challenge agent in the form of a liquid suspension after fasting for at least 90 minutes. The challenge procedure comprises two components: (i) a sodium bicarbonate solution (2.6g/120ml) pre-treatment for gastric acid neutralisation, followedby (ii) a 30ml 0.9% saline solution-bacterial challenge agent suspension.

We next evaluated the impact of sodium bicarbonate pre-treatment on the expected pH of the stomach at the time of bacterial challenge. Gastric acid represents a key defence mechanism against gastrointestinal infection. *Salmonella enterica* have evolved multiple strategies to improve survival following acid exposure, including the acid-tolerant response (ATR) and the arginine D-carboxylase system, but the organism still remains susceptible to gastric acid killing [32–34].

To model the gastric passage of the liquid challenge agent, we used the Rosset-Rice (RR) *in vitro* system to model bacterial stability and survival following simulated transit through the stomach and intestinal tract. The RR model is designed to replicate the low pH conditions of a fasted adult stomach by introducing acid at rates that mimic physiological acid secretion. The model is predicated on a set of assumptions concerning gastric physiology. In particular, the rate of gastric emptying is known to be regulated by several factors (25). In fasted adults, it is estimated that 240ml volume of non-caloric liquid can empty from the stomach in approximately 15-20 minutes [35,36]. The mean residual volume in a stomach after an overnight fast is estimated to range between 25-50ml [35,37,38]. Allowing for inter-individual variability, the resting pH of a fasted adult stomach typically ranges from pH 1 to 3 [39]. The basal acid output in healthy adult stomachs is estimated to be 1-5 mmol HCl/hour, which increases significantly on eating [40]. Although it mimics some key aspects of gastric physiology, one limitation is that it does not simulate the removal of gastric contents which would usually dilute the acidic environment as contents are removed over time.

Gastric pH dynamics were modelled in the modified Rossett-Rice *in vitro* system modelling an antacid dose of 2.6g sodium bicarbonate buffer to ensure that it maintains pH above the critical threshold of pH 3.5 identified for D23580 and 4/74 survival. Serial measurements every minute confirmed that gastric pH is maintained above the critical threshold pH >3.5 and is approximately pH 6.5 at 70 minutes following administration of bicarbonate buffer **(Figure 2 – Panel A)**.

**Figure 2.**
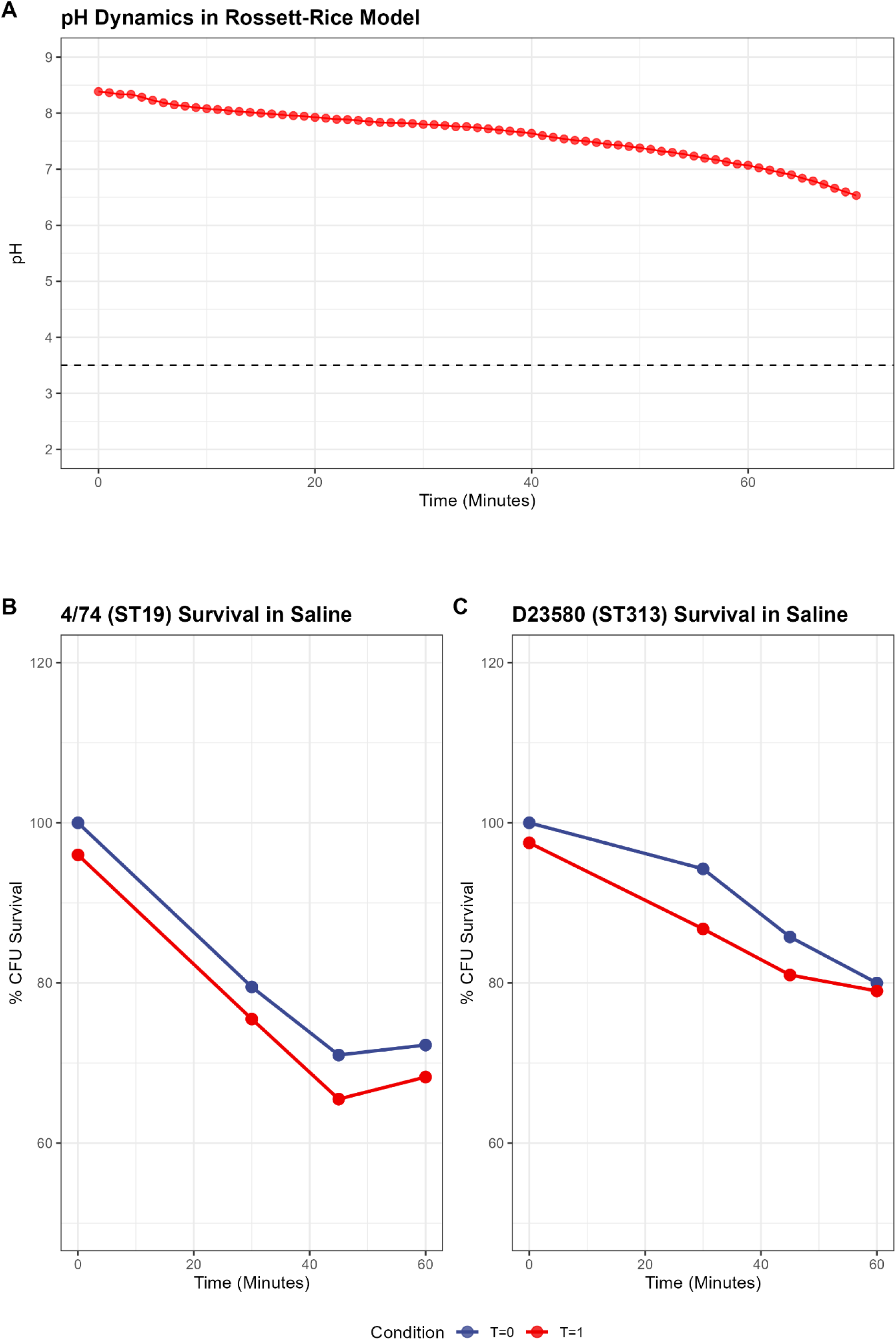
Simulated gastric fluid pH and D23580 and 4/74 survival in the modified Rossett-Rice *in vitro* system. (A) Simulated gastric fluid pH dynamics in the RR model following administration of 0.26g bicarbonate. A peristaltic pump (set at a rate of 0.4 ml per minute) was used to add additional SGF throughout the experiment to mimic conditions in the stomach, with pH monitoring every minute. pH >6.5 is maintained to 70 minutes. (B) 4/74 and (C) D23580 survival in the modified Rossett-Rice in vitro system, simulating bacterial transit through the upper gastrointestinal tract following T=0/T=1hr pre-incubation in saline. Mean % CFU survival calculated from 4 technical replicates per time point.

**Figure 3.**
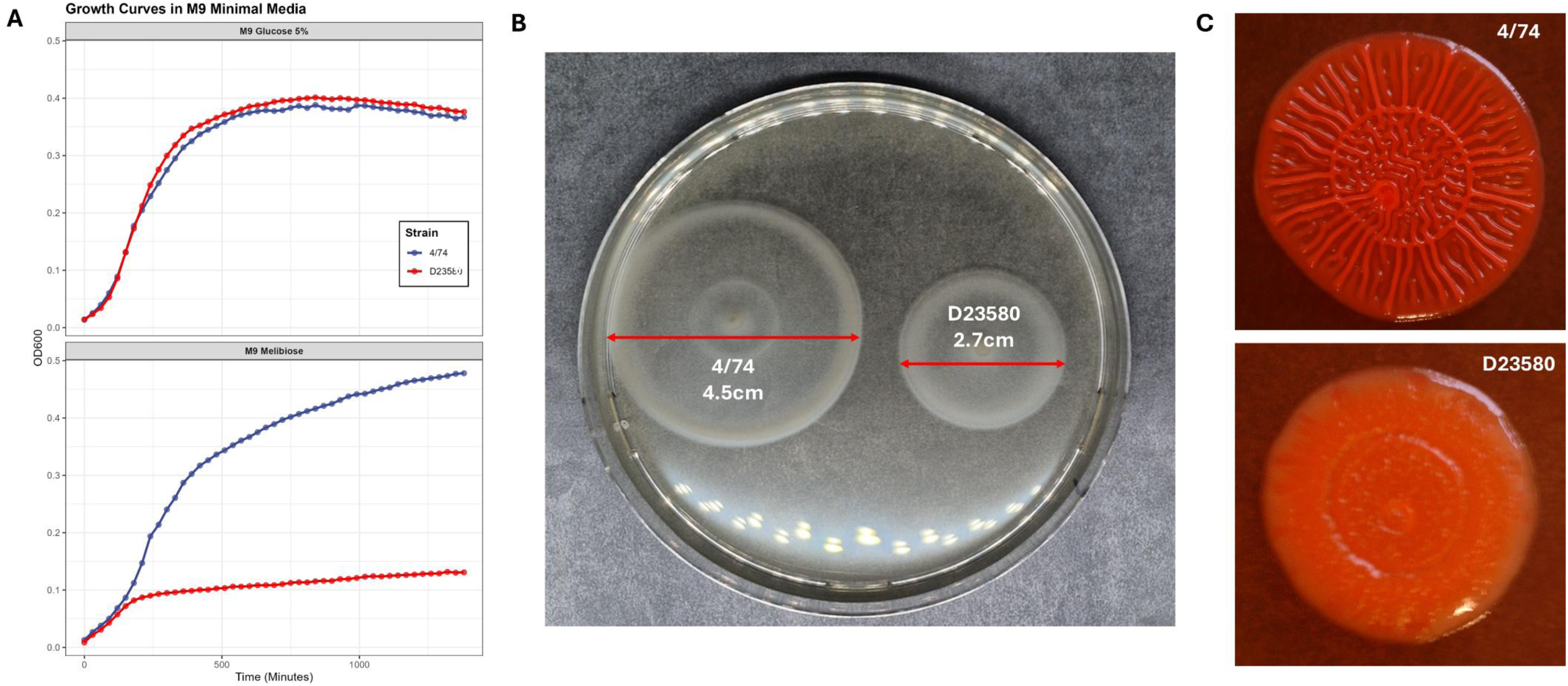
Phenotypic characterisation of STm 4/74 and D23580. (A) Growth curves for 4/74 and D23580 in M9 minimal media supplemented with 5% glucose (top) and melibiose (bottom). Analysed by Enzyscreen Growth Profiler over 24 hours, starting OD600 = 0.01. Comparable growth patterns are observed in 5% glucose, whilst growth of D23580 is inhibited in melibiose. Mean OD600 values at each time point calculated from six technical replicates. (B) Swimming motility of 4/74 (left) and D23580 (right) in 0.3% LB agar. Photographed after 4 hours incubation at 37°C. (C) Red, dry and rough (RDAR) morphotype colonies of (B) 4/74 and (C) D23580 after 72 hours incubation at room temperature on NaCl-deficient LB agar supplemented with Congo red. Imaged using the ImageQuant 4000 biomolecular imager.

### 3.6 *S.* Typhimurium viability remains stable in simulated gastric and intestinal fluids after 1 hour hold on ice

The modified Rossett-Rice model was then utilised to determine bacterial stability and survival following simulated transit through the gastrointestinal tract. Challenge strains were incubated in 0.9% saline solution or 2.6g/120ml sodium bicarbonate and either directly added to the RR model or after a 1- or 2-hour hold on ice. Fluid was then sampled at 0, 30, 45 and 60 minutes after addition to the RR model and plated for overnight incubation. Initial testing with D23580 demonstrated significant reductionin survival following T=2 hour on ice compared with T=0 hours across all sampled time points (p<0.05) (**Supplementary Figure 1**). Subsequent testing evaluated survival of both strains following T=0hr or T=1hr hold in ice in 0.9% saline. There were no significant differences in survival between T=0hr and T=1hr hold on ice prior to addition to the RR model. CFU counts declined with time and overall survival was approximately 70% and 80% after 60 mins in the RR model for 4/74 and D23580, respectively (**Figure 2** – **Panels B-C**).

We further modelled survival of the challenge strains during transit from the stomach to the small intestine. STm challenge strains were again subjected to the RR model and after 70 minutes samples were taken and exposed to simulated intestinal fluid and subsequently plated at 0, 30, 70-minute time points to mimic residence time in the upper small intestine. There was no difference in survival of either challenge strain between 0-70 minutes of exposure to simulated intestinal fluid (**Supplementary Figure 2**).

### 3.7 Phenotypic Confirmation and Characterisation

#### 3.7.1 STm D23580 and 4/74 exhibit differential growth characteristics in melibiose

A key metabolic property distinguishing D23580 from 4/74 is the differential ability to utilise disaccharide melibiose as a sole carbon source [28]. D23580 contains three nonsynonymous SNPs in melibiose utilisation genes (*melB* and *melR*) leading to downregulated expression of proteins responsible for active transport of melibiose across the cell membrane. It is speculated that the ability of 4/74 to metabolise melibiose confers a survival advantage during gastrointestinal colonisation, and the loss of this capacity in D23580 reflects niche adaptation towards an invasive phenotype [28]. This differential utilisation was demonstrated by evaluation of bacterial growth curves in minimal media and LB. Growth curves were comparable between strains in minimal media supplemented with 5% glucose, whilst growth of STm D23580 is markedly attenuated in minimal media supplemented with melibiose (**Error! Reference source not found.** – **Panel A**).

#### 3.7.2 STm D23580 and 4/74 exhibit differential swimming motility

As previously described, swimming motility in soft agar is reduced in D23580compared with 4/74 due to downregulation of flagellin expression [28]. This phenotype is consistent with other invasive *Salmonella* serovars including Typhi, Paratyphi A, and Gallinarum, which rapidly reduce flagellin expression upon cellular invasion to avoid immune recognition by TLR-5 and flagellin-specific CD4^+^ T-cells [41–43]. The reduced swimming motility of D23580 (halo diameter 2.7cm) compared with 4/74 (4.5cm) in soft agar after 4 hours incubation at 37°C is illustrated in **Error! Reference source not found. – Panel B**.

#### 3.7.3 STm D23580 and 4/74 exhibit differential RDAR morphotypes

At room temperature, many strains of *Salmonella* can form red, dry, and rough (RDAR) colonies, which result from the formation of aggregative fimbriae. RDAR colonies reflect multicellular behaviour that enhances *Salmonella* environmental stress resistance and biofilm formation. This loss of multicellular behaviour has been demonstrated in D23580, hypothesised to occur as a consequence of human host-adaptation [29]. A positive RDAR phenotype appears as a red colony with filamentous texture, whilst a negative RDAR phenotype appears as a smooth and yellow colony. Differential RDAR morphotype appearances between 4/74 and D23580 are illustrated in Error! Reference source not found. **– Panel C**.

#### 3.7.4 Antibiotic Susceptibility

Multi-drug resistance in *Salmonella spp.* is defined as resistance to antibiotics from three or more separate classes [44]. iNTS is increasingly multi-drug resistant in Africa, demonstrated by the emergence and persistence of ST313 [5]. D23580 harbours four plasmids, including pSLT-BT, which encodes resistance to chloramphenicol, ampicillin, streptomycin, sulphonamides, and trimethoprim.

Disc diffusion on Mueller-Hinton agar, and broth microdilution using BD Phoenix provided zone inhibition diameters and MIC concentrations respectively and were compared to published EUCAST guidelines for sensitivity [27]. Antibiotic sensitivity results for both strains were as predicted in both disc diffusion and broth microdilution methods. BD Phoenix minimum inhibitory concentrations (MIC) and interpretations are illustrated in **Table 2**.

**Table 2:** Antibiotic susceptibility profiles of STm 4/74 and D23580 as measured by broth microdilution using the automated BD Phoenix system. Minimum inhibitory concentrations (MIC) were measured and interpreted in accordance with EUCAST guidelines to determine susceptibility. S = susceptible, R = resistant

| Antimicrobial | MIC | Interpretation |
| --- | --- | --- |
| **4/74 (ST19)** |  |  |
| Ampicillin | ≤2 | **S** |
| Amoxicillin-Clavulanate | ≤2/2 | **S** |
| Ceftriaxone | ≤0.5 | **S** |
| Ciprofloxacin | ≤0.25 | **S** |
| Gentamicin | ≤1 | **S** |
| Meropenem | ≤0.125 | **S** |
| Piperacillin-Tazobactam | ≤4/4 | **S** |
| Trimethoprim-Sulphamethoxazole | ≤1/19 | **S** |
| **D23580 (ST313)** |  |  |
| Ampicillin | >8 | **R** |
| Amoxicillin-Clavulanate | 16/2 | **R** |
| Ceftriaxone | ≤0.5 | **S** |
| Ciprofloxacin | ≤0.25 | **S** |
| Gentamicin | 2 | **S** |
| Meropenem | ≤0.125 | **S** |
| Piperacillin-Tazobactam | ≤4/4 | **S** |
| Trimethoprim-Sulphamethoxazole | >4/76 | **R** |

### 3.8 Whole genome sequencing confirms STm genomic stability post-manufacture

Illumina whole-genome sequencing was performed to assess for single nucleotide polymorphism (SNP) differences which may have arisen during the manufacturing process compared with established reference strain sequences.

No SNP mutations were detected comparing *S*. Typhimurium 4/74 or D23580 manufactured stocks with the reference strain sequences [4/74 Genbank Accession Numbers: CP002487.1 - CP002490.1; D23580 Genbank Accession Numbers: LS997973.1 – LS997977.1], highlighting overall genomic stability. Consistency with genetic sequences from reference strains ensures the manufactured stocks administered to participants encode the same repertoire of virulence factors as circulating strains causing iNTS in sSA. Moreover, these strains encode the putative key antigens targeted by vaccine candidates in development, including the *wbaV, wbaU, wbaN, wzxB1* genes in the O-antigen biosynthesis cluster encoding O4, alongside *fliC* and *fliB* genes encoding phase variable flagellar antigens.

### 3.9 Sodium bicarbonate exposure does not impact key STm virulence gene expression

Administration of sodium bicarbonate pre-treatment is frequently utilised in enteric pathogen CHIMs to lower the required challenge inoculum, and to reduce batch-to-batch variation [24,45,46]. We have demonstrated that STm challenge strain survival is stable, and gastric pH is maintained above critical thresholds in a simulated gastrointestinal tract. However, sodium bicarbonate exposure has also been shown to have a significant effect on bacterial virulence factor gene expression in other enteric pathogens, notably in *E. coli* spp, *Citrobacter,* and *V. cholerae* [47–50]. We investigated the impact of sodium bicarbonate exposure on *S.* Typhimurium virulence gene expression; in case it altered the virulence of challenge agent stocks at the time of oral administration.

Transcriptomic profiling of STm D23580 and 4/74 has previously been undertaken and characterised in multiple infection-relevant conditions [28]. In this investigation, we performed RNA-seq on whole RNA extracted from STm D23580 and 4/74 after 30-minutes incubation in sodium bicarbonate solution (0.53g/30ml) and compared gene expression levels to cultures which had not been exposed to bicarbonate. These conditions were selected for testing to reflect the anticipated duration bacterial cells wouldbe in contact with bicarbonate buffer during transit from the stomach to the small intestine following oral challenge.

The transcriptional profiles of both strains were evaluated and individual gene expression level calculated as transcripts per million (TPM). A gene is considered as expressed when TPM is greater than 10 [28]. The proportion of expressed genes decreased after incubation in sodium bicarbonate solution for both 4/74 and D23580 (4.11% for 4/74 and 8.15% for D23580). Both strains expressed approximately 70% genes (TPM >10) after 30 minutes incubation in sodium bicarbonate solution compared with water (**Supplementary Figure 3**).

*Salmonella* pathogenicity islands SPI-1 and SPI-2 play major roles in human infection [51,52], with SPI-1 being required for bacterial invasion of epithelial cells and SPI-2 mediating the formation and maintenance of *Salmonella*-containing vacuoles (SCVs) [53]. Flagella are also major virulence factors during *Salmonella* infection [54]. It has been shown that sodium bicarbonate exposure down-regulates flagella expression in *E. coli* [50]. To determine the impact of sodium bicarbonate exposure on *Salmonella* virulence gene expression, we used a differential expression approach to compare gene expression before and after sodium bicarbonate treatment and determined that 6% of 4/74 genes and 5% of D23580 genes were differentially expressed. No significant up-regulation or down-regulation of SPI-1, SPI-2 or flagellar genes was detected in either strain following bicarbonate exposure. To determine any likely functional impact of the bicarbonate-regulated genes, we examined gene ontology and determined that there were no common functional links among the genes (**Figure 4**). Our data suggest that the minor changes in gene expression that were observed are more likely to reflect experimental variability than a meaningful biological trend. In summary, the transcriptomic data suggest that exposure to sodium bicarbonate does not significantly alter *Salmonella* virulence.

**Figure 4.**
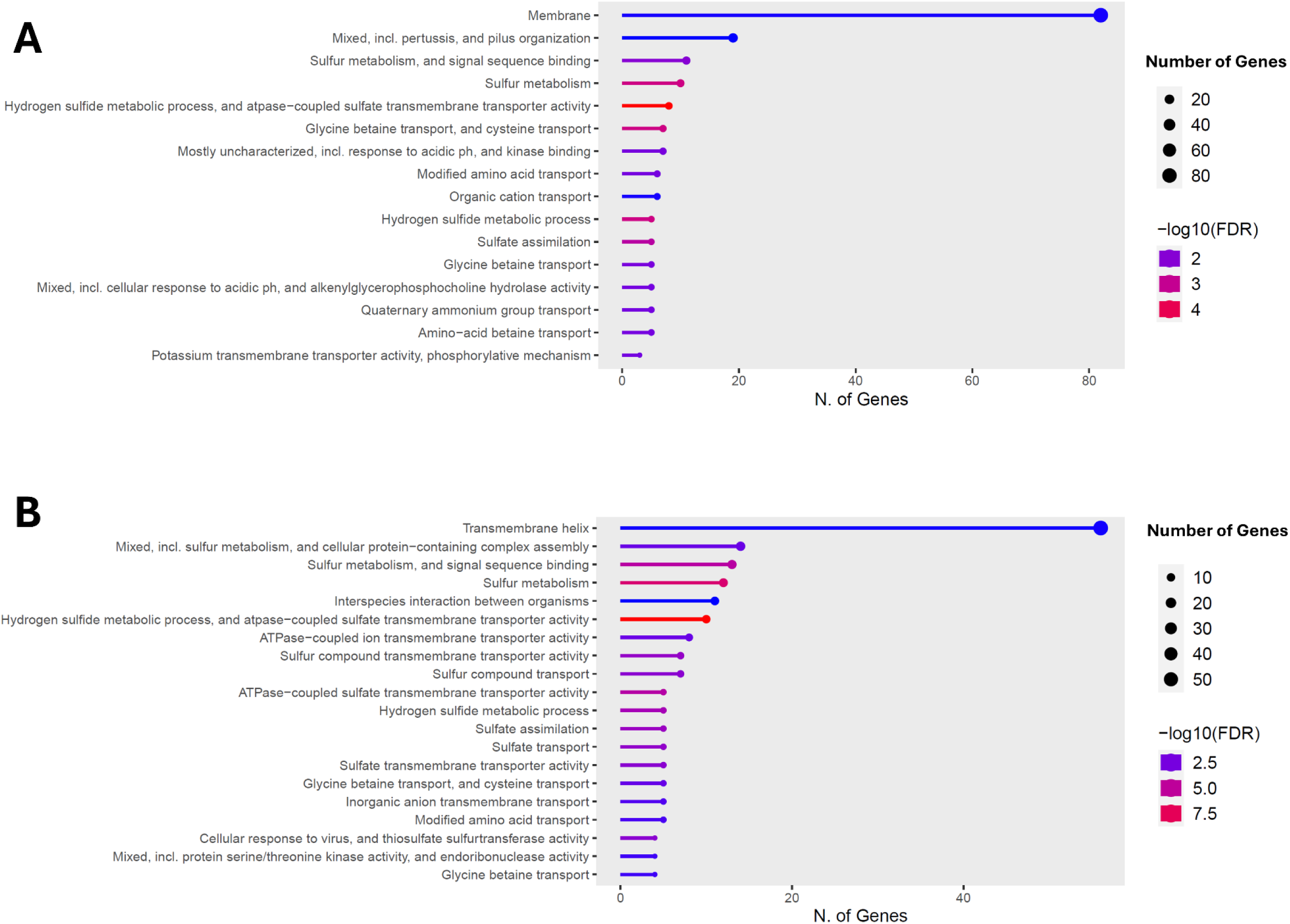
(A) STm 4/74 (ST19) and (B) D23580 (ST313) functional ontology of genes that were significantly differentially expressed following exposure to bicarbonate. Membrane and transmembrane helix associated genes were affected with greatest frequency; no differentially expressed genes were associated with SPI-1, SPI-2 or flagellar virulence genes.

## 4 Discussion and Conclusions

Bacterial stocks used for controlled human infection studies need to be manufactured to a high standard. In this study, we have described the processes involved with GMP manufacture and comprehensive phenotypic, genomic and transcriptomic characterisation of two *S.* Typhimurium strains - D23580 (ST313) and 4/74 (ST19). The steps described here are comparable with those undertaken for previous *Salmonella* challenge studies and serve as a benchmark for the type of testing required for new challenge agent development. Phenotypic and genotypic characterisation demonstrated that these two strains behaved as expected and no significant changes were introduced during the manufacturing process, giving us confidence prior to their administration to healthy human volunteers. We identified that both strains demonstrated improved survival in 0.9% saline holding buffer compared with sodium bicarbonate, subsequently adapting our challenge protocols in response to these data, modifying approaches used in previous *Salmonella* challenge models. A novel aspect of this study is a more detailed characterisation of *Salmonella* challenge stocks under conditions mimicking those encountered in the gastrointestinal tract using a modified Rossett-Rice model. The addition of a transcriptomic assessment of virulence gene expression offers a snapshot of the expected transcriptional state of the challenge agent as it enters the stomach, providing new insights and assurances that key virulence factors are unchanged in *Salmonella* following sodium bicarbonate pre-treatment which is a requirement of accepted enteric challenge model protocols.

The strainsused in this model were selected with careful consideration of the principles outlined in **Table 1**. An ideal challenge model would incorporate multiple diverse strains to better reflect pathogen diversity. Whilst this approach might be optimal, it is generally acknowledged that practical limitations, including GMP manufacturing costs and regulatory complexity, often preclude challenge studies from using more than one strain. Consequently, a detailed characterisation of well-documented strains is considered the most feasible approach in most circumstances. Provenance, traceable history from isolation, and prior comprehensive characterisation provides greater assurance in the safety and reliability of this first-in-human CHIM. *S.* Typhimurium was chosen as the preferred serovar for use in this model as it is responsible for the greatest burden of iNTS in sSA and is supported by the most comprehensive *in vitro* characterisation data.

Utilising two distinct strains of *S.* Typhimurium offers a unique opportunity to evaluate the clinical, microbiological and immunological outcomes following challenge in a healthy immune competent population. This platform facilitates direct comparison between strains and therefore offers a deeper understanding of the possible impact of genome degradation and human-adaptation observed in D23580, which may partly account for its propensity to cause invasive disease in sSA [6]. This also supports our approach to using wild-type D23580 which harbours a distinct plasmid and prophage repertoire compared with 4/74. Whilst this strain is classified as multi-drug resistant, it remains fully sensitive to all clinically relevant antibiotics used in the CHANTS study, ensuring participant safety.

In this study, challenge agent manufacture has been undertaken to GMP standard. At the time of writing, the UK does not legally require challenge agents to be produced under GMP. Approval from the UK Medicines and Healthcare products Regulatory Agency (MHRA) is only necessary if the CHIM study involves an investigational medicinal product, such as if a vaccine is administered alongside the challenge agent. Nevertheless, the MHRA guidance generally recommends following GMP principles to ensure the safety and quality of the produced challenge agent. Oversight primarily rests with the ethics committee and study sponsors [55,56].

*Salmonella* Typhimurium is often ascribed as one of the most comprehensively studied and characterised pathogens and is a model organism for bacteriological research. Availability of such a compendium of characteristic data was leveraged to facilitate the comprehensive phenotypic and genomic characterisation undertaken in this study [57]. *In vitro* phenotypic properties were confirmed to be consistent with published data including RDAR morphotype, swimming motility and growth kinetics in a variety of nutrient media. Antimicrobial susceptibility testing also confirmed activity of antibiotic agents selected for use within the CHANTS study. Further, whole genome sequencing has confirmed genomic stability of both strains throughout the manufacturing process and confirmed no genetic modifications which might alter expected pathogenicity. Longitudinal sequencing of stored challenge agent stocks is also being undertaken throughout the duration of the CHANTS study to confirm maintained genomic stability.

Building on work from previous *Salmonella* challenge models, we have undertaken novel assessments of the stability and viability of *Salmonella* in a range of pH conditions, holding buffers, and in a simulated gastrointestinal environment, to model survival and phenotype at the point of oral administration.

Our data indicated differences in the stability of *S.* Typhimurium in holding buffers compared with previous *S.* Typhi and Paratyphi studies. Recent typhoidal *Salmonella* challenge studies administer 2.1g/120ml of sodium bicarbonate buffer followed by the challenge agent in a 0.53g/30ml sodium bicarbonate holding buffer [45]. The data presented here indicate improved bacterial survival in a 0.9% saline holding buffer compared with sodium bicarbonate. As a result, the sample preparation SOP was updated so that the *S.* Typhimurium challenge agent is administered in 30ml 0.9% saline, similar to the holding buffer used in recent *Shigella sonnei* challenge studies [58]. In parallel, the bicarbonate buffer protocol was adjusted to account for the reduction in sodium bicarbonate that would have been otherwise used as a holding buffer for the 30ml *S.* Typhimurium inoculum. We now administer 2.6g/120ml sodium bicarbonate solution, which provides the same total mass of bicarbonate ensuring equivalent buffering capacity compared to previous *Salmonella* challenge study SOPs [45].

Bacterial cells remained stable in pH >3.5, stable for up to 2 hours in 0.9% saline holding buffer and 1 hour in sodium bicarbonate and remained stable up to 60 minutes in a modified Rossett-Rice model. These key findings provide assurance that viable bacterial cell counts remain stable during the challenge agent preparation, transfer and administration protocols followed in the CHANTS study. Furthermore, we have demonstrated that exposure to bicarbonate buffer does not significantly alter *Salmonella* virulence factor gene expression. These findings taken together provide assurance that bacterial cell concentration remains stable during challenge agent preparation and transfer from the laboratory to clinic, and that gastric acid neutralisation with bicarbonate pre-treatment does not impact pathogenicity at the point of oral administration, thus ensuring close recapitulation of what likely occurs in natural exposure.

Whilst this study has provided a comprehensive characterisation of STm 4/74 and D23580 for use in a first-in-human CHIM, there are limitations in our approach which should be considered. Firstly, contemporary epidemiological genomic and surveillance studies highlight the important contributions *S.* Enteritidis – and the emergence of *S.* Typhimurium ST313 sub lineages 2.2 and 2.3 – have in iNTS disease burden [5,59]. Including these strains in our model could further enhance generalisability to those isolates currently circulating in sSA, which account for a significant proportion of iNTS disease. Furthermore, using recently harvested isolates – or attenuated strains – could potentially compromise the rigorous ethical standards adhered to in the development of CHIMs, as highlighted in **Table 1**. As such, strain selection was based upon these core principles, with STm 4/74 and D23580 being selected as priority pathogens which could provide the greatest translational data within the context of a CHIM.

In developing a non-typhoidal *Salmonella* (NTS) challenge model to mimic invasive disease, we opted to develop two strains of *S.* Typhimurium for administration to volunteers, as the serovar responsible for the majority of iNTS cases globally. Future plans include the development of a *Salmonella* Enteritidis challenge model, which accounts for approximately one third of iNTS cases worldwide. Like *S.* Typhimurium, *S.* Enteritidis strains associated with invasive disease in low-income settings, including *S.* Enteritidis ST11 [60], have distinct genetic profiles compared with those causing gastrointestinal disease in high-income regions. The strain selection, development and manufacture of such a strain would follow principles similar to those outlined here.

In summary, extensive characterisation of both *S.* Typhimurium D23580 and 4/74 has experimentally confirmed predicted and known properties of these strains. These investigations have significantly advanced the development of *Salmonella* human infection models and support ongoing efforts to gain deeper understanding of host-pathogen interactions and immune correlates of protection. Gaining such new understanding of iNTS infection can strengthen ongoing efforts to support iNTS vaccine and therapeutic development. We aim to report CHANTS study primary outcome data in the near future and will outline how these findings could be translated to support ongoing efforts to combat iNTS disease in sSA.

## 5 Declarations

### Ethics Statement

This CHANTS study protocol has been reviewed and approved by the NHS Health Research Authority (London—Fulham Research Ethics Committee 21/PR/0051; IRAS 301659) and is registered on ClinicalTrials.gov (registration number NCT05870150). Study sponsor Imperial College London.

### Funding

This work is funded by The Wellcome Trust (grant number 224029/Z/21/Z: awarded to principal investigator MMG). CS, GSC, and MMG are supported, in part, by the NIHR Imperial Biomedical Research Centre. LL is funded by an EMBO fellowship (ALTF 95-2023). MAG is funded by an AXA Research Professorship. The funders had no role in study design, data collection and analysis, decision to publish, or preparation of the manuscript.

### Availability of Data and Materials

The datasets used and/or analysed during the current study are available from the corresponding author on reasonable request.

Genbank Accession Numbers (Reference Genomes):

*Salmonella* Typhimurium 4/74 CP002487.1 - CP002490.1

*Salmonella* Typhimurium D23580 LS997973.1 – LS997977.1

Raw RNA-seq data from bicarbonate experiments have been submitted to the NCBI Gene Expression Omnibus (GEO) Repository. Accession codes are currently being generated, are available from the corresponding authors, and will be included in a final published manuscript.

### Competing Interests

The authors have declared that no competing interests exist.

### Author Contributions

Conceptualisation – CS, MAG, GSC, RKMC, JCDH, MMG

Data Curation – CS, AB, XJ, JAW, NR, AES

Formal Analysis – CS, AB, XJ, LL

Funding Acquisition – MMG

Investigation – CS, AB, XJ, JAW, NR, AES, AR, RV, ZK, BPS

Project Administration – ES

Supervision – GSC, RKMC, JCDH, MMG

Writing – Original Draft Preparation – CS, AB, JAW, XJ

Writing – Review and Editing – All Authors

## 8 Supplementary Figures

**Supplementary Figure 1:**
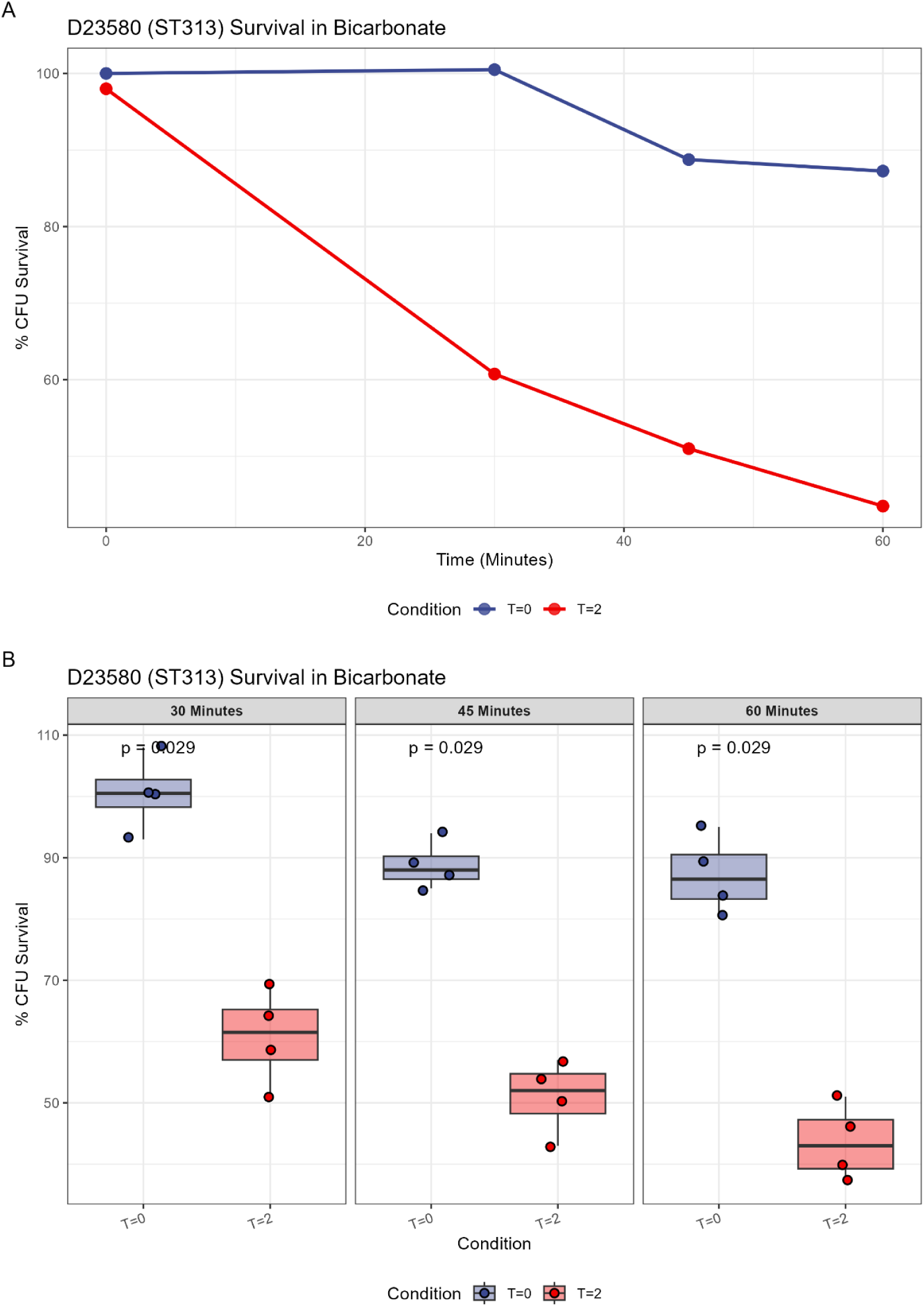
(A) and (B) represent D23580 survival over time following incubation in 2.6g/120ml sodium bicarbonate solution for either T=0hr or T=2hr prior to addition to the RR model. Boxplots represent median and interquartile range. P = two-sided Mann-Whitney U test.

**Supplementary Figure 2:**
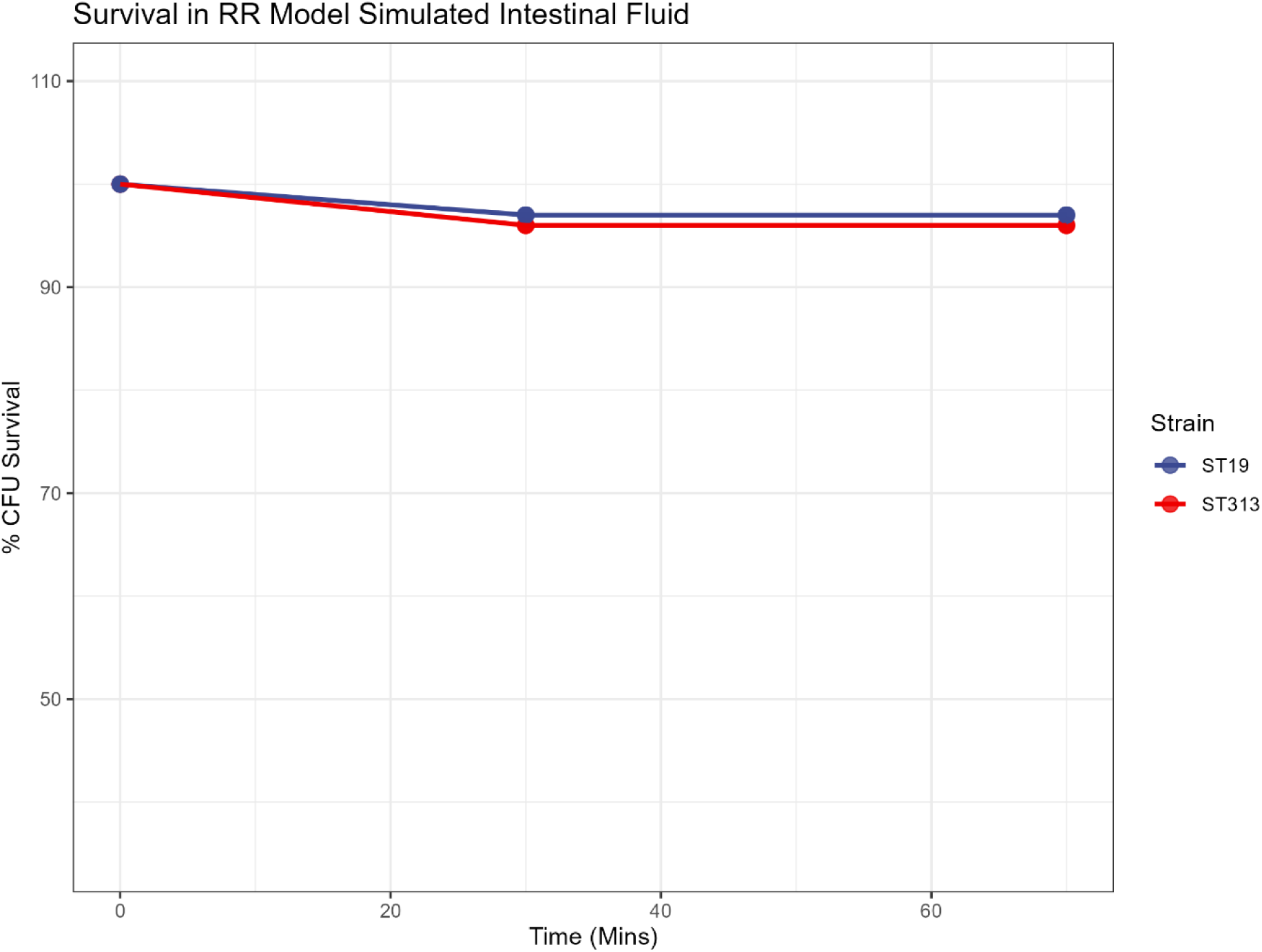
Percentage CFU survival of 4/74 (ST19) and D23580 (ST313) following incubation in simulated intestinal fluid to model passage through the upper gastrointestinal tract.

**Supplementary Figure 3:**
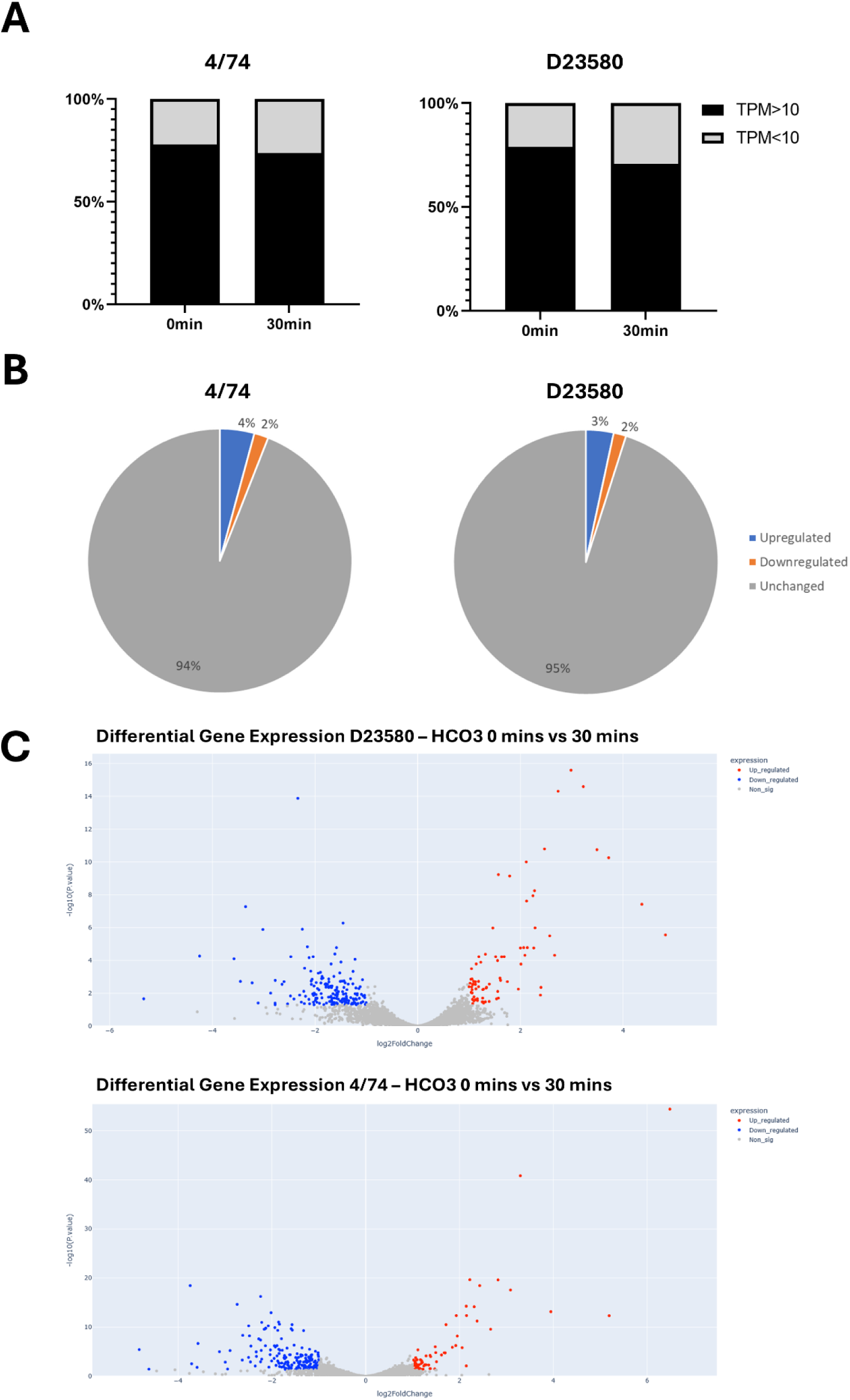
(A) Relative gene expression levels of 4/74 (left) and D23580 (right) in TPM. (B) Proportion of significantly up- or down-regulated genes in 4/74 (left) and D23580 (right) after 30-minutes exposure to sodium bicarbonate. (C) Volcano plots representing differentially expressed genes. Interactive plots with gene label names are available at https://github.com/Adalijuanluo/CHANTS_RNAseq.

**Supplementary Figure 4:**
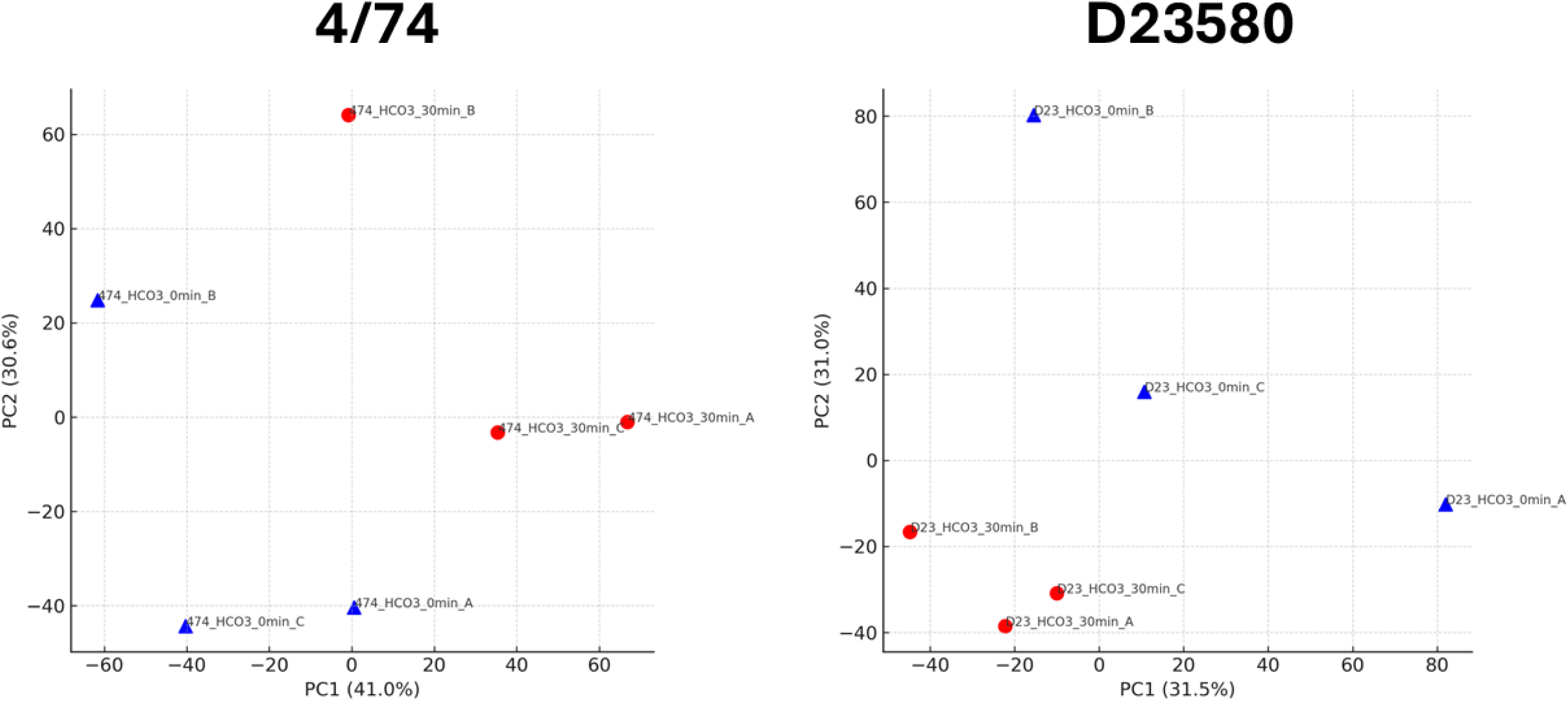
Principal component analyses representing variance in gene expression patterns in 4/74 (left) and D23580 (right) after 0-minutes or 30-minutes incubation in sodium bicarbonate solution.

## Notes

https://github.com/Adalijuanluo/CHANTS_RNAseq

### Competing Interest Statement

The authors have declared no competing interest.

### Summary of Updates

In this revised manuscript, we have expanded the Methods section to include additional procedural detail on GMP manufacture, microbial limits testing, the Rossett-Rice model, and RNA extraction and sequencing. The Transcriptomics section has been updated with further description of the RNA-seq analysis and confirmation of data deposition in the NCBI Gene Expression Omnibus (GEO). Figures have also been reformatted into multi-panel layouts to enhance clarity and presentation of the data.

